# Costs and benefits of mating with fertilized females in a species with first male sperm precedence

**DOI:** 10.1101/386045

**Authors:** Leonor R Rodrigues, Alexandre RT Figueiredo, Thomas Van Leeuwen, Isabelle Olivieri, Sara Magalhães

## Abstract

Different patterns of sperm precedence are expected to result in specific mating costs and benefits for each sex, generating different selection pressures on males and females. However, most studies concern species with mixed paternity or last male sperm precedence, neglecting species with first male sperm precedence, in which only the first mating is effective.

Here, we measured costs and benefits of multiple mating for both sexes of the spider mite *Tetranychus urticae*. First, we assessed the stability of the sperm precedence pattern, by mating females to one, two or several males, immediately after the first mating or 24 hours later. We found complete first male precedence, independently of the mating interval and the number of matings. Females paid a cost of polyandry, as multiply-mated females laid fewer eggs than once-mated females. However, while first males had reduced survival when exposed to an intermediate number of virgin females, second males paid no additional costs by matings with several mated females. Moreover, by mating multiply with mated females, males decreased the total number of offspring sired by first males, which suggests that these matings may entail a relative benefit for second males, despite being ineffective.

Our results show that complex costs and benefits may arise in males in species with first male precedence. How these costs and benefits affect the maintenance of selection for polyandry remains an open question.

## Introduction

Multiple matings are prevalent in internally fertilizing species (Simmons, 2001). However, this behaviour is expected to entail negative consequences for both sexes, such as increased risk of predation and energy expenditure (G Arnqvist & Nilsson, 2000; Göran Arnqvist & Rowe, 2005). Consequently, for multiple mating to be selected, the reproductive advantage gained by this behaviour should to be superior to the costs incurred.

Polyandry, i.e., multiple mating in females with different males within a breading season (G Arnqvist & Nilsson, 2000; Taylor, Price, & Wedell, 2014), is expected to be selected whenever females obtain direct benefits, such as increased fecundity and survival with each mating via nuptial gifts or nutritious ejaculates. For instance, in the bruchid beetle *Callosobruchus maculatus*, substances in the seminal fluids lead to an increase in the number of offspring produced (Eady, Wilson, & Jackson, 2000). Alternatively, but not exclusively, females can mate multiply to increase the quality or diversity of their offspring, in which case their benefit is indirect (Kvarnemo & Simmons, 2013; Snook, 2014). This has been observed, for example, in the bumble bee *Bombus terrestris*, in which polyandry can increase colony resistance to parasites by maximizing the chances that at least some individuals survive (Schmid-Hempel & Baer, 1999).

From a male’s perspective, the benefits of multiple mating are quite straightforward, assuming that more matings result in more offspring production. Yet, polyandry, an expected outcome of increased male mating rate, can result in the offspring of a single female being shared by several males. As a result, males have evolved adaptations to sperm competition, such as harassment, altered male genitalia and toxic ejaculates, so as to sire a higher share of offspring of each female (Simmons, 2001). This, in turn, can be costly for females, decreasing their fitness (e.g., Chapman et al. 1995). Consequently, the balance between costs and benefits obtained with each mating is probably not the same for both sexes (Bateman, 1948). This imbalance can give rise to sexual conflicts, in which one sex employs reproductive tactics to enforce matings while the other resists them, depending on which sex will benefit the most from each mating (Göran Arnqvist & Rowe, 2005).

From the interaction between adaptations that have evolved in males and females to increase their reproductive success, emerges a pattern of sperm precedence. Such patterns may be more beneficial for one sex than for the other, thereby generating different selection pressures on each sex. In species with mixed sperm precedence, in which paternity is shared by several males, selection in males should favour increased mating frequency as a result of offensive and defensive adaptations to sperm competition (Mark Ridley, 1989) but for females the benefits obtained from each additional mating are not as straightforward (Göran Arnqvist & Rowe, 2005; Bateman, 1948). Indeed, females may obtain genetic benefits from a more diverse or fit offspring, but may pay the cost of excessive matings, which can lead to sexual conflict over mating rate. However, under complete first male precedence, in which only the first mating of a female is effective, genetic benefits cannot be obtained. Thus, in the absence of direct benefits, selection in these species should favour monandry, as both sexes are expected to invest all resources in copulations involving virgin females only (Thomas, 2011), thereby limiting the scope for sexual conflict (Hosken, Stockley, Tregenza, & Wedell, 2009). However, monandry may be imposed by males on females, as a result of evolved defensive adaptations against sperm competition. Once monandry is achieved, selection for this enforcement may be relaxed, potentially leading to polyandry being restored. In addition, females may be selected to gain the opportunity to choose the sperm they will use to sire their offspring (Dougherty, Simmons, & Shuker, 2016), and second males may be selected to obtain a share of the females’ offspring (offensive traits). For instance, it has been suggested that second males may increase their relative reproductive success from matings with mated females without obtaining any paternity share, by displacing or killing the sperm of the first male inside the female (Harshman and Prout 1994, Macke et al. 2012). All these patterns suggest that the sperm precedence observed in a species is not necessarily evolutionarily stable (Dougherty et al., 2016). Furthermore, in several species, the pattern of sperm precedence has been shown to vary according to multiple ecological factors, such as the number of matings (Zeh & Zeh, 1994), the interval between mating events (Bullini, Coluzzi, & Bianchi Bullini, 1976), the effectiveness of the first mating (Weldingh, Toft, & Larsen, 2011), or differences in male traits (*e.g*., size) (Bissoondath & Wiklund, 1997).

Because of the lability of this pattern, it is important to know (a) how it translates into cost and benefits for both sexes and (b) if it is maintained under different ecological conditions. Nevertheless, most of the research has been done in species with mixed or last male sperm precedence (Simmons, 2001) and empirical studies addressing costs and benefits for both sexes in species with first male precedence are remarkably scarce (but see Fisher et al. 2013; Boulton and Shuker 2015, 2016).

Here, we study the consequences of mating for both sexes in the two-spotted spider mite, *Tetranychus urticae.* Earlier studies suggest that in this species only the first copulation of a female is effective (Helle, 1967). This leads to the expectation that males should only mate with virgin females to avoid unnecessary costs. Accordingly, males actively guard juvenile quiescent females and mating occurs as soon as females moult into virgin adults (Potter, Wrensch, & Johnston, 1976), a behaviour that is consistent across species with first male sperm precedence (M. Ridley, 1989). In addition, when given the choice between mated and virgin females, males, prefer to mate with virgins, basing their decision upon volatiles and chemical trails (Oku, 2010; Rodrigues, Figueiredo, Varela, Olivieri, & Magalhães, 2017). Nevertheless, mated females are often observed mating (Clemente, Rodrigues, Ponce, Varela, & Magalhães, 2016; Oku, 2010). Here, we provide a comprehensive account of potential costs and benefits of polyandry for both sexes in spider mites. First, we performed paternity tests, using a recessive mutation that codes for resistance to a pesticide as a genetic marker, in order to describe the pattern of sperm precedence. To account for the ecological lability of the pattern sperm precedence, we varied the number of matings and the interval between mating events. We then measured the fecundity and survival of females that re-mated at different time points, to assess potential costs or benefits of polyandry for females. In addition, we analysed differences in male survival in the presence of different numbers of virgin or mated females to assess potential costs of multiple mating for first and second males, respectively. Finally, we compared the total number of offspring sired by first males mated to females with different mating status, to assess the potential benefits for second males.

## Materials and Methods

### Spider mite populations, rearing conditions

To study the costs of multiple mating in males, we used a population collected in Carregado, Portugal, established at the University of Lisbon in 2010 from approximately 300 individuals (TuTOM). To study the other traits, we used the EtoxR strain, resistant to etoxazole (Van Leeuwen et al., 2012), and the LondonS strain (Grbić et al., 2011), susceptible to the same pesticide, both established at the University of Lisbon in 2013 from approximately 2000 individuals. Etoxazole is a pesticide that interferes with chitin synthesis, affecting spider mite embryos and juvenile stages at the time of hatching or ecdysis (i.e., at the quiescent stage; Van Leeuwen et al. 2012). In the EtoxR strain, resistance to Etoxazole is recessive and conferred by a single chitin synthase 1 (CHS1) amino acid change (Van Leeuwen et al., 2012). We used pesticide resistance as a marker for paternity.

All populations followed an antibiotic treatment to eliminate symbionts, using a protocol adapted from Breeuwer (1997). Prior to the experiment, we confirmed that resistance was fixed in the EtoxR strain and absent in the LondonS strain, following a protocol adapted from Van Leeuwen et al. (2012). We compared the fitness of females mated with resistant or susceptible males, to account for potential male genotype effects, and no significant differences were found (Fig. S1).

All spider-mite populations were reared in large numbers (>2000) on bean plants (*Phaseolus vulgaris*, Fabaceae, var. *Enana*; Germisem Sementes Lda, Oliveira do Hospital, Portugal), under controlled conditions (25°C, photoperiod of 16L: 8D).

### Experimental Setup

#### Mating protocol

Randomly selected virgin females from the EtoxR strain were allowed to mate once, twice or multiple times (O, T and M, respectively) and the mating interval between the first and subsequent matings was either 0 or 24 hours (re-mated immediately, I or re-mated later, L, respectively; Fig. 1a). In treatments with two or more matings, females mated either first with a resistant and then with susceptible males, or the opposite. Thus, these females were allocated to 5 different treatments: O, TI, MI, TL and ML. A description of these treatments is provided in Figure 1a. Briefly, EtoxR quiescent females were isolated for 24 hours on leaf discs on water-saturated cotton without males. Once they became adult (one day later), groups of 5 females were placed with 5/6 susceptible - or resistant - males on 0.8 cm^2^ leaf discs. The patches were observed for 2 hours and once a female had successfully mated, it was transferred to a new patch, either empty or with males of the other strain, in the same proportion (5 females: 6 males). Half of the females placed with males on the second patches were observed for two more hours and isolated when mated (TI). The other half was left unobserved on the patch with males for 24 hours (MI), which, in this species, is a sufficiently large time interval to ensure the occurrence of multiple matings (authors Pers. Obs., Krainacker and Carey 1989; Magalhães et al. 2007). The females left alone after the first mating on the first day were either left alone for one more day (O) or transferred to patches with males of the alternative strain. Again, half the females were observed for 2 more hours and isolated if mated (TL) and the other half was left unobserved on the patch with males for 24 hours, thus allowing for multiple matings (ML). After mating, two-to three-days old females were isolated on a 2.55 cm2 leaf disc placed on water-soaked cotton, in order to measure the life-history traits mentioned below. To maximize the effectiveness of the first mating, we (a) isolated males prior to testing them, (b) limited their copulations to 5 females, a value below their daily reproductive limit (Krainacker & Carey, 1989) and (c) discarded first matings interrupted by other individuals on the patch. Due to excessive experimental effort, this experiment was done on 22 separate days, all treatments being represented at each day.

**Figure 1.**
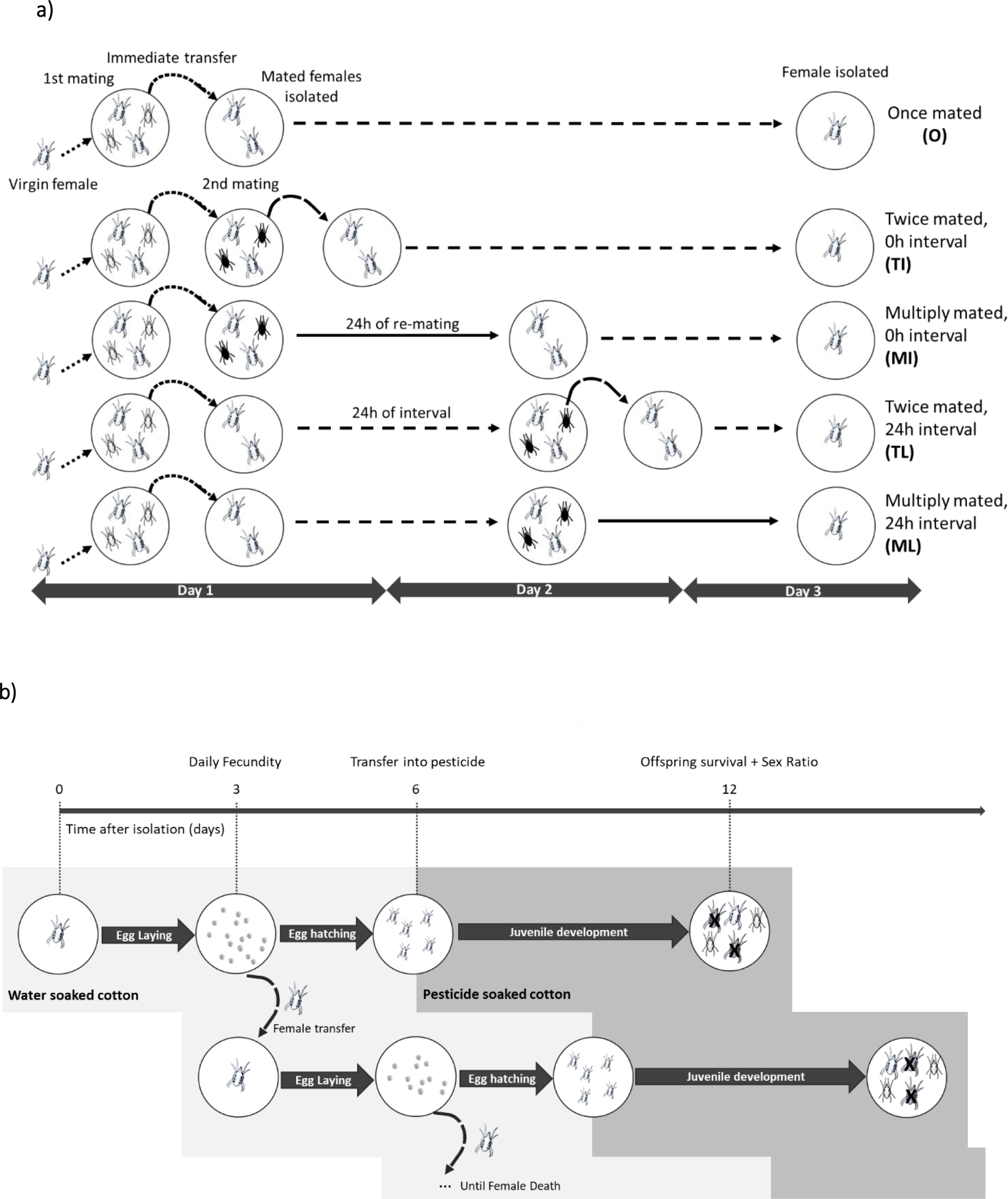
Protocol followed to assess sperm precedence and female fecundity and survival. **a) Mating Protocol.** Groups of 5 females and 5/6 males were placed together on patches until they mated. Females mated once (O), twice (T) or multiply (M) with a mating interval between the first and subsequent matings of either 0 hours (I, Immediately) or 24 hours (L, later). Females are bigger than males and are always resistant to pesticide (white). Males are smaller and represented in black or white. Different male colours represent different phenotypes after pesticide exposure: black males are susceptible, while white males are resistant to pesticide. Note, that in this scheme only one male type order is represented (resistant males first). However, both orders were performed. Dotted arrows: females were transferred immediately from one patch to the next; full arrows: females were maintained on a patch for 24 hours with males; dashed arrows: females were maintained on a patch for 24 hours without males. **b) Data Collection.** Each female was isolated on a leaf disc placed in water-soaked cotton and its survival was checked daily. Every 3 days, the female was transferred to a new leaf disc where she could continue to lay eggs. The number of eggs laid by the female on each leaf disc was measured after each transfer. On the 6th day, the leaf disc was moved into a container with cotton soaked in diluted pesticide. On the 12th day offspring sex-ratio and survival were measured, to extract offspring paternity and the total number of daughters sired by the first male. X, death owing to pesticide.

#### Data Collection

##### Effect of multiple mating on offspring number and paternity

We tested whether the number of matings and the interval between matings affected offspring number, indicating potential costs or benefits of mating multiple times for females. Moreover, we assessed the paternity of that offspring, as a measure of potential benefits for first or second males. For this purpose, females isolated on leaf discs were transferred every three days to a new leaf and the eggs laid on the old leaves were counted. Daily fecundity was measured as the total number of eggs laid per female divided by the number of days the female was alive (Fig. 1b). To assess paternity, the eggs laid by the isolated females were allowed to develop until they reached the first juvenile stage (three days after female transfer to a new disc), then leaf discs were transferred to water-soaked cotton with diluted etoxazole (500 ppm). Six days later, the number of adult daughters, adult sons and dead juveniles on each leaf disc was recorded (Fig. 1b). Spider mites are haplodiploid, producing haploid sons, which result from unfertilized eggs, and diploid daughters, stemming from fertilized eggs (Helle & Sabelis, 1985). Therefore, the number of alive daughters and dead juveniles indicate, respectively, the number of offspring sired by resistant and by susceptible males. The survival of sons should not be affected by pesticide application since all females used were resistant to etoxazole and sons only inherit the genetic material of their mothers. Note however, that natural death in the quiescent stage may be confounded with death by pesticide exposure. Yet, because this occurs in all treatments, including the once-mated treatment that serves as control, the differences between treatments are a true measure of paternity share, excluding natural death at the quiescent stage. In total, we analysed the daily fecundity of 485 females and assessed the paternity of offspring from 377 females.

##### Effect of multiple mating on male and female survival

To determine whether mating multiply benefited females by increasing their survival, we tested whether female survival varied with the number of matings and the interval between matings. To this aim, the same females used to assess paternity and daily fecundity were used to measure female survival (Fig. 1b). The survival of mated females was followed daily after female isolation on a 2.55 cm2 leaf disc placed on water-soaked cotton. In total, we analysed the survival of 485 females.

A different experiment was performed to measure the costs of mating in terms of survival in first and second males. To this aim, males and females were isolated separately at the quiescent stage, to control their age and ensure virginity prior to the experiment. When these individuals became adults (*circa* 24 hours later), groups of ten females were either left isolated (virgin – V) or placed with 15 males (mated – M). The latter were left with the males for 24 hours to ensure the occurrence of multiple matings (authors Pers. Obs., Krainacker and Carey 1989; Magalhães et al. 2007). The next day, focal virgin one-day old males were placed on a leaf circle with the previously isolated females: Males were allocated to leaves with either 1, 5 or 20 virgin females (V1, V5, V20), which allowed testing costs for first males, or with 1, 5 or 20 mated females (M1, M5, M20), thereby testing for potential costs in second males. To normalize densities across treatments, patch size varied according to the number of individuals (0.38 cm^2^, 2.55 cm^2^ or 9.1 cm^2^ for patches receiving 1, 5 or 20 females, respectively). The focal male was then transferred daily to a new patch with the same number of (mated or virgin) females in every treatment except for the ones with 20 females. In this last treatment, as male mating capacity decreases with age (Krainacker & Carey, 1989), from the third day onwards, the focal male was placed with 12, instead of 20 females (size of the patch: 6.25cm^2^). Every day until death, male survival was recorded. In total, the survival of 180 males was analysed. Due to excessive experimental effort, and the very high number of females required for each replicate, this experiment was carried out on 66 separate days.

##### Potential benefits of ineffective matings for males

Mating with mated females may provide a relative increase in the fitness of second males, despite first male sperm precedence. For example, by mating with mated females, males may displace or kill the sperm inside the female and thereby increase their relative reproductive success (Macke et al. 2012). Here, we tested whether multiple matings could reduce the genetic contribution of first males. The total number of daughters (i.e. male genetic contribution to the next generation) sired by first males mated to females with different number of matings and mating intervals, was compared. The same females used to assess paternity and daily fecundity were used to measure this trait (Fig. 1b). Because the aim was to study lifetime fecundity, females who died due to artificial causes (drowning in water-soaked cotton) were excluded from the analysis. In total, we analysed the total number of daughters produced by 427 females.

### Statistical analyses

All analyses were carried out using the R statistical package (v. 3.0.3). Maximal models were simplified by sequentially eliminating non-significant terms from the highest-to the simplest-order interaction, with the highest p-value to establish a minimal model (Crawley 2007; see Table S1), and the significance of the explanatory variables was established using chi-squared tests, in the case of discrete distributions or Wald F tests, in the case of continuous distributions (Bolker et al. 2008; see Table S2). *A posteriori* contrasts with Bonferroni corrections were done to interpret the significant effect of factors with more than two levels (glht, multcomp package): comparisons were done between treatments with single matings, or single females in the case of male survival, and all other treatments (Table S3).

To analyse the effects of mating on female survival, daily fecundity, total number of daughters and offspring paternity, the mating treatment (i.e. 0: once-mated, TI: twice-mated immediately, MI: multiply-mated immediately, TL: twice-mated later, ML: multiply-mated later) was fit as fixed explanatory variable, whereas day and male type order (female mated first with a resistant and then with susceptible males, or the opposite) were fit as random explanatory variables.

To analyse the effects of mating on male survival, the female status (i.e., M: mated; V: virgin) and the number of females on each patch (1, 5, 20) were fit as fixed explanatory variables, and day was fit as a random explanatory variable.

#### Effect of multiple mating on offspring number and paternity

To analyse the proportion of offspring sired by the first male, we redistributed the data of offspring survival into two variables called contribution of the 1st male (1M) and contribution of the second male (2M) to offspring. 1M corresponds to the number of dead juveniles or the number of alive daughters, depending on whether the first male was susceptible or resistant, 2M corresponds to the number of alive daughters or the number of dead juveniles, depending on whether the first male was susceptible or resistant. These parameters were computed using the function cbind, with 1M, 2M and the number of sons as arguments. Since the model was greatly over-dispersed, we used a generalized linear mixed model with a beta-binomial error distribution and added the term ziformula=~1 to the model (glmmTMB, glmmTMB package) (Brooks et al., 2017).

Daily fecundity per female was transformed to improve normality (Box-Cox transformation; Crawley 2007) and subsequently analysed using linear mixed-effect models (lmer, lme4 package).

#### Effect of multiple mating on male and female survival

Male and female survival were analysed using a Cox proportional hazards mixed-effect models (coxme, coxme package). In the analysis of male survival (MS), because the interaction between the fixed factors was significant, we analysed separately each level of female status for the effect of female number.

#### Potential benefits of ineffective matings for males

The total number of daughters sired by the first male was analysed using the variable “contribution of the 1st male” (1M). This parameter was analysed using a model with negative binomial distribution (glmer.nb, lme4 package) to account for data overdispersion.

## Results

### Effect of multiple mating on offspring number and paternity

Overall, there was no significant effect of the mating treatment (Χ^2^_4_=1.411, P=0.842) on the proportion of offspring sired by the first males (Fig. 2a). Therefore, first male sperm precedence is virtually complete. This also indicates no differences in sex-ratio across treatments. However, mating treatment affected daily fecundity significantly (F_4,389.95_=8.633, P<0.001). Contrast analyses revealed that females that mated multiple times 24h after their first mating had significantly lower fecundity compared to once-mated females, while females from all other treatments laid the same number of eggs (O vs TI: Z=-0.025, P =1.00, O vs TL: Z=-0.725-, P=1.00, O vs MI: Z = −1.976, P = 0.193 and O vs ML: Z=-4.151, P<0.001; Fig. 2b, Table S3).

**Figure 2.**
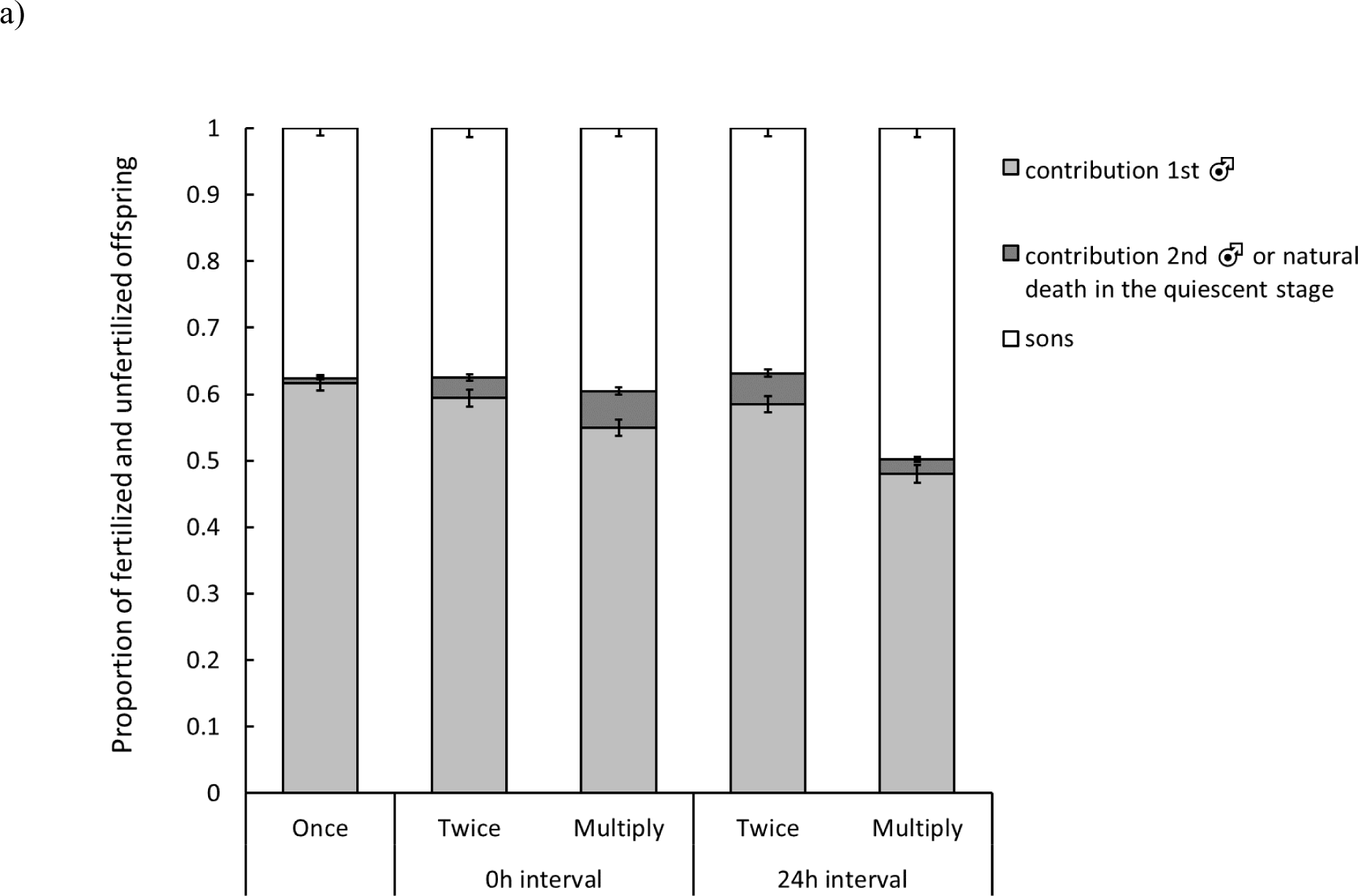
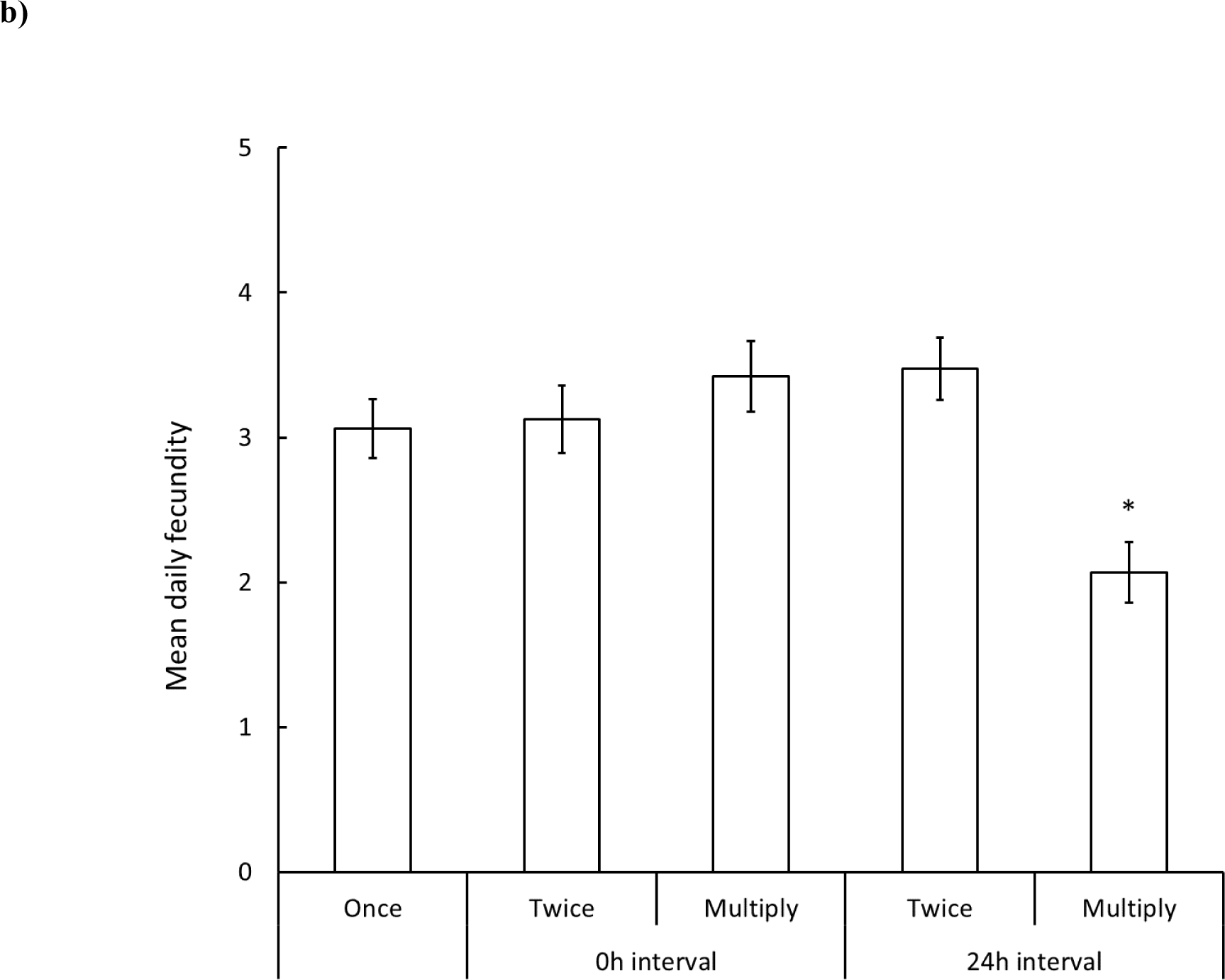
Effect of multiple mating on offspring number and paternity. Females mated once (O), twice (T), or multiply (M). Re-mating was set immediately (0h interval; I) or 24 hours after the first mating (24h interval; L). **a) Proportion of fertilized and unfertilized offspring across treatments.** Fertilized offspring (i.e. daughters) is divided into proportion of daughters sired by the first (light grey) and by the second male (dark grey). Note however, that natural death in the quiescent stage may be confounded with death by pesticide exposure, in both bars representing fertilized offspring. Unfertilized offspring (sons) is represented in white. Vertical bars correspond to standard errors of the mean. **b) Mean number of eggs laid daily by females.** Vertical bars correspond to standard errors of the mean. Asterisk (*) represent significant level (P < 0.05).

### Effect of multiple mating on male and female survival

The mating treatment the female was subjected to affected significantly the survival of females (X^2^_4_=10.899, P=0.0277). However, no significant differences were found when comparing all treatments to the once-mated control (O vs TI: Z=-0.203, P =1.00, O vs TL: Z=1.235, P=0.867, O vs MI: Z = 1.379, P = 0.671 and O vs ML: Z=-1.719, P=0.343; Fig. 3a). As for males, their survival was significantly affected by the interaction between female status and the number of females on each patch (X^2^_2_=7.198, P=0.027). Indeed, males placed with virgin females survived less in the presence of 5 females than in presence of 1 female per day (V1 vs V5: Z=2.349, P=0.038; V1 vs V20: Z=0.353, P=1.00; Fig. 3b, Table S3). However, no significant differences in survival were observed when males were placed with mated females (X^2^_2_=0.497, P=0.78; Fig. 3c).

**Figure 3.**
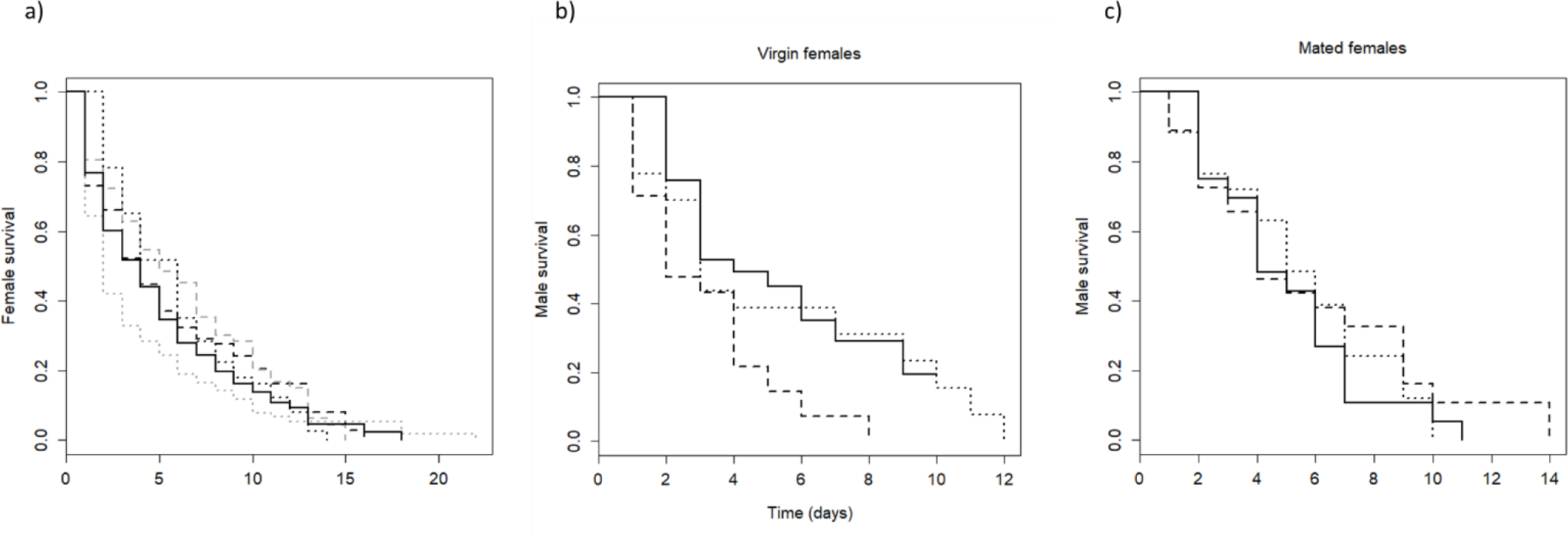
Effect of multiple mating on male and female survival. **a) Female survival curve.** Females mated once (O), twice (T), or multiply (M). Re-mating was set immediately (0h interval; I) or 24 hours after the first mating (24h interval; L). Each line corresponds to different number of matings: continuous line, one mating; dashed line, two matings; dotted line, multiple matings. Grey lines correspond to L rematings; black dashed and black dotted lines correspond to I re-matings. **b) and c) Male survival curves.** Males were placed in patches with 1, 5 or 20 virgin (b) or mated (c) females every day and its survival was followed. Distinct types of lines represent different number of females per patch status: continuous line, one female; dashed line, 5 females; dotted line, 20 females.

### Potential benefits of ineffective matings for males

A significant effect of mating treatment was found for the total number of fertilized offspring sired by the first male (X^2^_4_= 15.956, P=0.003). Indeed, multiply-mated females with an interval of 24 hours between first and subsequent matings produced fewer fertilized offspring, compared to once mated females (O vs ML: Z= 3.174, P= 0.006; Fig. 4, Table S3). This suggests that second males benefit by mating with mated females. However, females belonging to all other treatments produced the same number of fertilized offspring than once-mated females (O vs TI: Z=-0.024, P=1.00, O vs TL: Z=-0.315, P=1.00; O vs MI: Z= −0.367, P=1.00; Fig. 4, Table S3).

**Figure 4.**
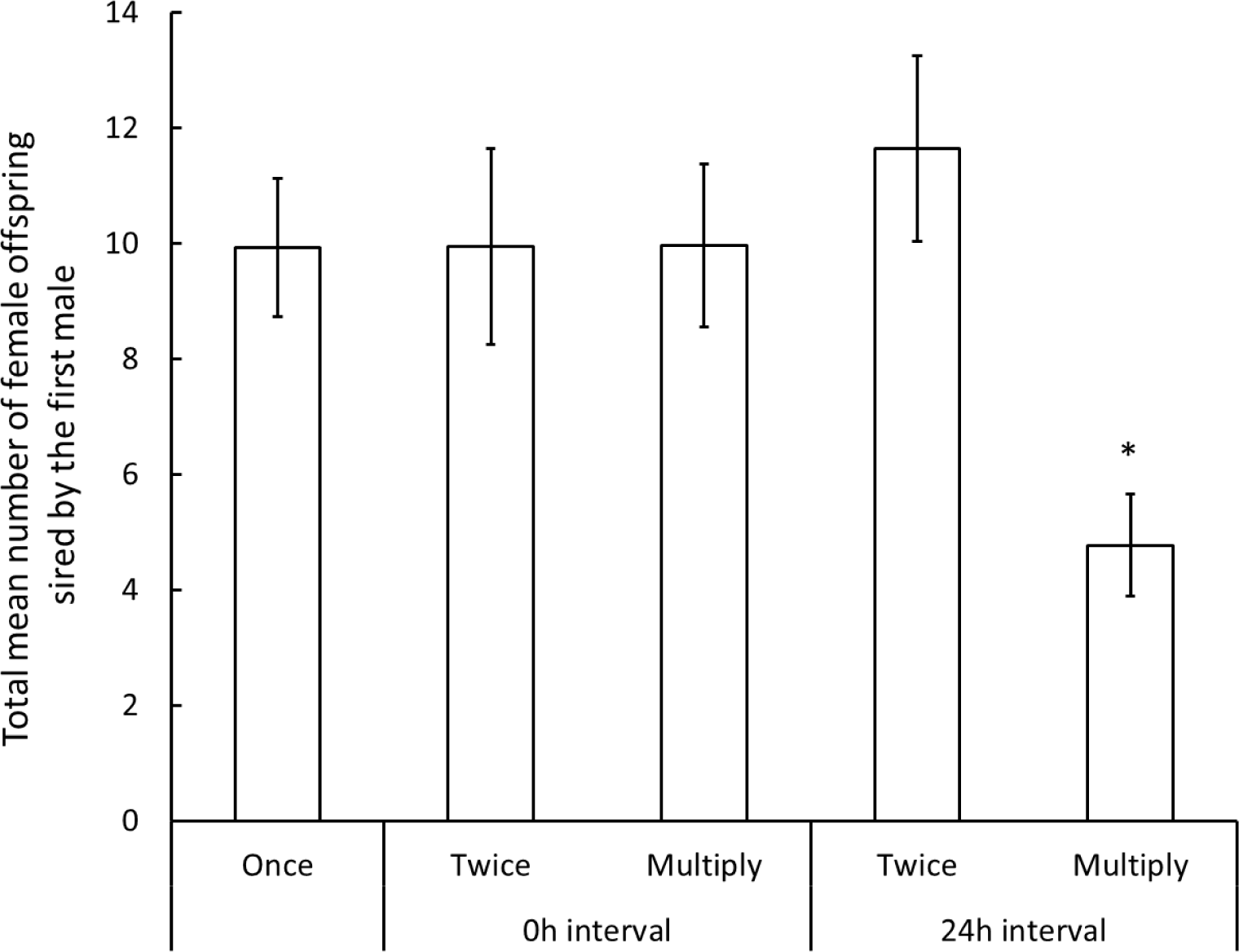
Total mean number of offspring sired by the first male. Females mated once (O), twice (T), or multiply (M). Re-mating was set immediately (0h interval; I) or 24 hours after the first mating (24h interval; L). Vertical bars correspond to standard errors of the mean. Asterisk (*) represent significant level (P < 0.05).

## Discussion

Our study revealed that nearly all fertilized offspring was sired by the first male, independently of the mating interval and the number of matings. In addition, a decrease in fecundity, but not in survival, was found in females that had multiple mating opportunities after an interval of 24 hours between the first and subsequent matings. Males, however, suffered increased costs of mating when placed with 5 virgin females daily, but not when placed with mated females, revealing costs for first, but not for second males. In addition, first males produced fewer daughters when the females they mated with re-mated with other males.

We cannot disentangle complete, to nearly complete, first male sperm precedence, because natural death in the quiescent stage may be confounded with death by pesticide exposure. Thus, there is a non-null threshold of detection for fertilization by second males. Still the contribution of second males to siring offspring in these conditions, if any, is extremely small, and is not likely to explain the existence of polyandry in this species. Some species with first male sperm precedence have been shown to change their pattern of sperm precedence with mating interval and number of matings. For instance, in the silk worm *Bombix mori*, the paternity share of the first male changes from 0.95 to 0.06 in two hours (Suzuki, Okuda, & Shinbo, 1996). However, other species, such as the wasp *Diadromus pulchellus*, keep their pattern of sperm precedence across mating intervals (Agoze, Poirié, & Périquet, 1995).

Still other factors, that could influence the pattern of sperm precedence remain to be tested. For instance, in *Drosophila pseudoobscura,* a mostly monandrous fly, females use the sperm from the second mating whenever the first mating opportunity failed (Fisher et al., 2013). The authors make a distinction between true polyandry and pseudopolyandry, that occurs when females remate but no sperm competition takes place, owing for instance to lack of sperm transfer. This could be the case in spider mites as well, a hypothesis that remains to be tested. Indeed, sperm depletion, or incomplete sperm transfer in the first male may allow for some paternity share between the first and second as sperm depletion is a phenomenon potentially common in spider mites (Krainacker & Carey, 1989). In addition, an early study, in which the effect of several mating intervals on sperm precedence in spider mites was tested without controlling for sperm depletion, found that second males can sire some offspring when the interval between copulations is shorter than 24 hours (Helle 1967). Unfortunately, the frequency of sperm-depleted matings in spider mite natural populations, which is expected to determine its role in shaping the evolution of sperm precedence, is unknown.

The fact that we could not detect evidence for first male sperm precedence being incomplete suggests that indirect genetic benefits of polyandry for females are absent in this species. Moreover, females that mated multiply paid a cost of fecundity. Most of the few studies that explored the costs and benefits of polyandry in species with first male sperm precedence show that male ejaculates provide benefits to the females (e.g. Thailayil et al. 2011; Helinski and Harrington 2012). However, in *Nasonia vitripennis*, a mostly monandrous wasp, female costs have been observed in patches with intense male harassment (Rebecca A. Boulton & Shuker, 2015). Since no differences in sex-ratio were observed between treatments, the mating costs observed here are, most likely, a reflection of the negative effects of multiple matings coupled with increased number of mating attempts, as observed in *N. vitripennis*. The fact that we only found a cost when the interval between the first and subsequent matings was of 24 hours, may be explained by differences in female receptivity across different mating intervals. Indeed, females that mated 24 hours after the first mating, independently of the number of matings, took longer to mate and interrupted matings more often than females that re-mated immediately after the first mating (authors personal observations, Clemente et al. 2016). This suggests that females become more resistant to mating sometime after the first copulation, consequently suffering increased costs.

In the absence of clear benefits of multiple mating for females and of direct benefits for males, if the costs associated with those matings are low, males may mate with mated females because there is no selection pressure to eliminate such behaviour. In several species, the cost of mating for males varies with the mating status of females. Matings with mated females may entail fewer costs, if males allocate sperm differently according to the reproductive value of females (“strategic ejaculates”; Simmons 2001; Kelly and Jennions 2011). Accordingly, in species with first male sperm precedence, we expect males to invest more in matings with virgin females, as those are the ones with the highest reproductive value. Our results are in line with this prediction, as they show that only first males (those that mate with virgins) pay a cost of mating. This suggests that males invest more in mating with these females, either increasing their mating rate or transferring more sperm in each copulation, although it is intriguing that fewer costs were detected at the highest female density. In contrast, second males, which mate with mated females, payed the same survival costs in patches with different female densities. Previous results show that copulations with virgin females occur at a faster rate and last longer than copulations with mated females (Rodrigues et al., 2017), which is in line with these results. Therefore, males may engage into matings with virgins, which result in high offspring yield but also a survival cost, or into matings with mated females, yielding no offspring but also fewer costs.

Despite being ineffective, matings with mated females may still yield some benefits to males. A decrease in the total fecundity of multiply-mated females has been observed in spider mites (Macke et al., 2012), a result that we recover here. This may translate into fewer offspring being sired by first males (the Relative Fitness hypothesis, Macke et al. 2012). Here, we validated this hypothesis by showing that first males produced fewer offspring (i.e., daughters) when mating with females that mated multiply 24 hours later. Because the proportion of daughters remained unchanged, this decrease in the number of daughters is probably due to a decrease in fecundity of females owing to costs of mating and male harassment. Therefore, mating with mated females can increase the relative reproductive success of subsequent males, by reducing the genetic contribution of the first males to the following generations. This strategy requires that the harming males (or their brothers) produce some descendance and pay a low penalty with this behaviour. Apart from the life-history costs of the behaviour, which we showed here to be low, they could lose mating opportunities with virgin females. Therefore, this behaviour should be most favoured in expanding populations: males can first mate with virgin females, then later, when these become scarce, turn to harming mated females, hence not suffering much from lost opportunities. Moreover, the uncovered benefits should be dependent on population structure. Indeed, in large populations, benefits should be mitigated as they are shared by all other males of the population, while In small populations, relatedness can be high, in which case, reducing the fitness of other, related, males in the population may not be advantageous (e.g., Carazo et al. 2014). Yet, this is contingent on the scale of competition, because if competition occurs locally, males are not expected to behave differentially towards related males, as these are their only competitors (Pizzari et al. 2015). Finally, the effectiveness of this behaviour relies on a collective action as one mating is not enough to reduce the fitness of the first. Therefore, the precise population structure in which this behaviour will be beneficial may be very specific. Roughly, it should correspond to a situation of budding dispersal (Gardner, Arce, & Alpedrinha, 2009), in which related males arrive together in a patch occupied by unrelated individuals, and collectively reduce the fitness of unrelated males for the next generation. Although spider mites are expected to face different conditions during their dispersal-colonization phases, it is not clear that such beneficial conditions occur often enough for this behaviour to be selected. Hence, the probability of selecting this trait in males will hinge on how often they will encounter the conditions favouring it. It is thus important to design experiments varying population structure to test these ideas.

Altogether, our results show that multiple mating is costly for females but that matings with mated females are potentially beneficial for males. The latter was not expected, given that we are dealing with a species with first male sperm precedence. The consequences of polyandry in such species have been seldom explored, leaving a gap in our knowledge (but see Dougherty et al. 2016). Indeed, if we had found no benefits of polyandry in both sexes, we could speculate that selection would be favouring monandry with time. Conversely, if males and females benefited with polyandry, we could expect that selection would maintain polyandry, which in turn, would open the door for an evolution of the sperm precedence pattern itself. Because we found that some males may benefit from mating with mated females, but that females suffer costs with polyandry, conflicts between sexes should be present and the direction of selection on polyandry will depend on which sex is winning this conflict.

## Acknowledgements

The authors wish to thank Inês Santos for help with the maintenance of mite cultures at cE3c and Emilie Macke, João Alpedrinha, Alison Duncan, Diogo Godinho and Suzanne Alonzo for insightful comments on the manuscript. These experiments were funded by Portuguese (FCT, Fundação para a Ciência e Tecnologia) and French (ANR, Agence Nationale de la Recherche) funds through an FCT-ANR project (FCT-ANR//BIA-EVF/0013/2012) to SM and IO. LRR had a Ph.D. Grant (SFRH/BD/87628/2012) funded by FCT. The authors have no conflict of interest to declare.

## Authors Contributions

Resources provisioning: SM, TVL; Experimental conception and design: LRR, SM; acquisition of data: LRR, ARTF; statistical analyses: LRR; paper writing: LRR, SM, with input from all authors. All authors have read and approved the final version of the manuscript.

